# Assisting walking balance using a bio-inspired exoskeleton controller

**DOI:** 10.1101/2022.10.19.512851

**Authors:** M. Afschrift, E. Van Asseldonk, M. Van Mierlo, C. Bayon, A. Keemink, H. van der Kooij, F. De Groote

## Abstract

Balance control is important for mobility, yet exoskeleton research has mainly focused on improving metabolic energy efficiency. Here we present a biomimetic exoskeleton controller that supports walking balance and reduces muscle activity. Humans restore balance after a perturbation by adjusting activity of the muscles actuating the ankle in proportion to deviations from steady-state center of mass kinematics. We designed a controller that mimics the neural control of steady-state walking and the balance recovery responses to perturbations. This controller uses both feedback from ankle kinematics in accordance with an existing model and feedback from the center of mass velocity. Control parameters were estimated by fitting the experimental relation between kinematics and ankle moments observed in humans that were walking while being perturbed by push and pull perturbations. This identified model was implemented on a bilateral ankle exoskeleton. The exoskeleton provided 30% of the estimated ankle moment during steady-state and perturbed walking. Across twelve subjects, exoskeleton support reduced calf muscle activity in steady-state walking by 19 % with respect to a minimal impedance controller. Proportional feedback of the center of mass velocity improved balance support after perturbation. Muscle activity is reduced in response to push and pull perturbations by 10 and 16 % and center of mass deviations by 9 and 18% with respect to the same controller without center of mass feedback. Our control approach implemented on bilateral ankle exoskeletons can thus effectively support steady-state walking and balance control and therefore has the potential to improve mobility in balance-impaired individuals.

## INTRODUCTION

Wearable robotic devices (e.g. exoskeletons and prostheses) are currently being developed to enhance mobility in able-bodied subjects, or to restore mobility in persons with musculoskeletal and neurological disorders. Recent developments in hardware and control enabled large reductions in muscle activity and metabolic energy consumption in able-bodied subjects during walking (*1–5*). Nevertheless, researchers still struggle to design controllers for wearable robotic devices that not only reduce effort, but also support balance control. The limited ability to support balance control has important negative implications as individuals with mobility impairments typically have reduced balance control resulting in a high fall incidence (*6–8*). Many potential users would potentially benefit from wearable robotic devices that simultaneously reduce effort and support balance during walking. Here, we developed and tested a biomimetic controller for a bilateral ankle exoskeleton that simultaneously supports steady-state and perturbed walking.

Commonly used exoskeleton controllers that prescribe kinematics or assistive moments are not easily extendable to include balance control as flexible balance control requires online feedback from the state of the human and exoskeleton. Traditionally, exoskeletons are controlled using pre-defined joint angle (*9*) or torque trajectories (*1*). Control approaches based on torque trajectories have been very successful in reducing metabolic cost in healthy users (*1–5*). As these control approaches do not provide balance support, balance impaired users have to use external stabilizers such as crutches or walkers that limit function (*7*). Two recent studies supported balance with a predefined torque trajectory that mimics the joint moment in response to perturbations in healthy subjects (*10*, *11*). These studies demonstrated the potential of exoskeletons to support balance. Yet, this control approach is not generalizable to different types of perturbations. Notably, the adaptability and stability of human locomotion originates from sensorimotor feedback.

Biomimetic exoskeleton controllers that are inspired by human reflex control during steady-state walking have been shown to be adaptable but they do not stabilize walking against whole-body perturbations. Geyer and Herr proposed a computational model of the musculoskeletal system that produces stable walking based on local feedback control (*12*). In this model, both muscle dynamics and local feedback provide stabilization against local perturbations. This model has been used to control exoskeletons and prostheses (*13–16*). In this case, the joint angles derived from encoders on the device are the input to the neuromuscular model whereas the joint moments are the outputs. The resulting controllers have been shown to be adaptable to different walking speeds (*17*). However, local feedback control is not sufficient to stabilize human movement against whole-body perturbations. In simulation, Song & Geyer showed that the neuromechanical model can predict the main changes in muscle activity in response to local perturbations (e.g. mechanical tap of tendons), but not to whole-body disturbances such as slip and trip perturbations (*18*). A neuromuscular controller for a transfemoral prosthesis leads to more robust walking in simulation compared to minimal impedance control, but does not capture human responses to mid-swing disturbances in hardware experiments (*19*). These results might not be surprising given that humans use sensory integration to shape feedback responses and thus do not rely solely on local feedback.

Supra-spinal feedback pathways have an important contribution in human standing and walking balance control (*20*). Changes in ankle muscle activity and moments after fore-afterward perturbations during standing and walking can be explained by delayed feedback of whole-body center of mass (COM) kinematics (*21–24*). The relation between muscle activity and delayed COM kinematics also indicates that supra-spinal mechanisms, and not only local feedback loops, are important in human balance (*20*, *25*). Using feedback control from COM kinematics might also improve the ability of wearable robotic devices to support balance control.

We developed an ankle-foot exoskeleton controller that aimed at assisting both steady-state and perturbed walking. Inspired by the observed relation between COM kinematics and reactive muscle activity in perturbation experiments in humans (*21*), the proposed exoskeleton controller relies on feedback of COM kinematics in addition to local feedback of ankle kinematics to estimate the required ankle moment. We identified control parameters in the underlying neuromechanical model by fitting simulated and measured ankle joint moments in a dataset with steady-state walking and walking with pull and push perturbations applied at the pelvis (*26*). The resulting neuromechanical model was used to control an ankle foot exoskeleton in a novel perturbation experiment. The exoskeleton provided 30% of the joint moment estimated by the model during steady-state and perturbed walking. Our main hypotheses were that additional feedback of COM kinematics is needed in a neuromechanical model to describe changes in joint moment after perturbations; and ankle-foot exoskeletons controlled with a neuromechanical model with additional feedback of COM kinematics will assist balance control. Given the interaction with the human, successful assistance might result in either reduced muscle activity and/or reduced deviations from steady-state locomotion in response to perturbations.

## RESULTS

### Parameter identification perturbed walking

We first identified control parameters of a neuromechanical model based on an existing motion-capture dataset. This dataset documents the response to pull and push perturbations applied at the pelvis at toe-off during slow walking at 0.62 m/s (*26*). We started from a state-of-the art neuromechanical model with virtual Hill-type muscles driven by local reflexes with ankle angles and ground reaction forces as input and joint moments as output (i.e. default neuromechanical model) (*12*). We extended this model using additional feedback of deviations in COM velocity with respect to steady-state walking. Control parameters were estimated by optimizing the fit between simulated and measured ankle moments during steady-state and perturbed walking for both the default neuromechanical model and the neuromechanical model with COM feedback.

Adding COM velocity feedback was needed to track the ankle moment in perturbed walking (Fig. 1). The root mean square error (RMSE) in ankle joint moment was 14 Nm in the default neuromechanical model and 9 Nm in the model with COM feedback (Fig 1. B-C). The model with COM feedback can predict the decrease in ankle moment in response to pelvis pull and the increase in ankle moment in response to pelvis push perturbations. In contrast, the default model predicts an increase in ankle moment, opposite to the observed decrease, in response to pelvis pull perturbations and no change in ankle moment in response to pelvis push perturbations (Fig. 1, D). Although we did not optimize the fit between measured and simulated muscle activity, the model with additional COM feedback also captured the increase in calf muscle activity in response to push perturbation and the increase in tibialis anterior activity in response to pull perturbations (appendix, Fig. S2).

**Fig. 1.**
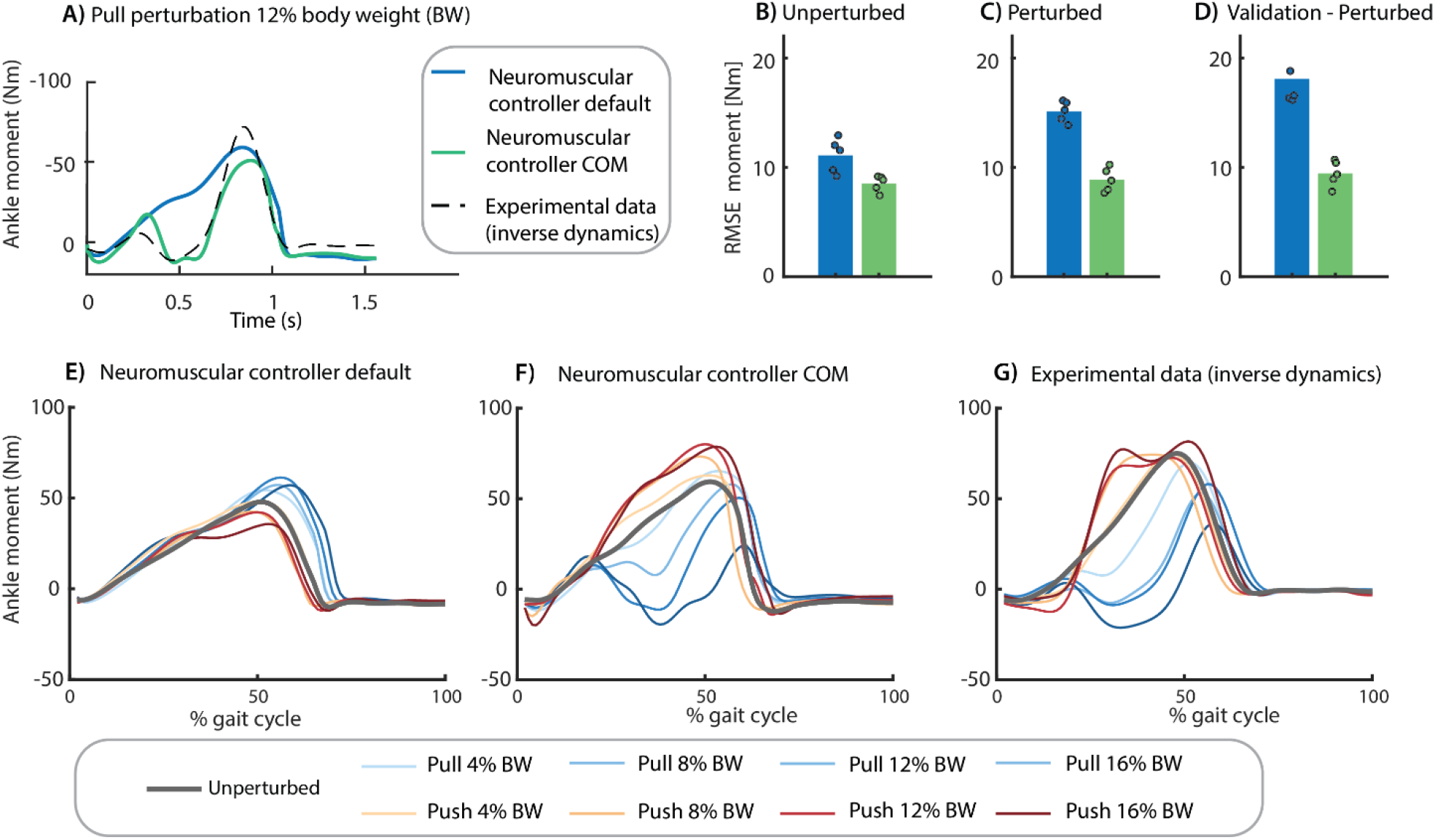
Control parameter identification. We estimated the control parameters of a model without (blue) and with (green) additional feedback of COM velocity by minimizing the difference between the experimental ankle moment and the ankle moment simulated with the neuromechanical model (representative example in Panel **A**). Control parameters of each model were estimated by tracking eight steady-state and 16 perturbed gait cycles in one optimization problem. Perturbations were applied at toe-off of the contralateral leg while the subjects walked at 0.62 m/s. The RMSE between experimental and simulated ankle moments was smaller in the model with additional COM feedback for **(B)** steady-state walking, **(C)** perturbed walking, and **(D)** a validation perturbation trial that was not used in the parameter estimation (the dots represent the RMSE in each of the five subjects and the bar represents the average across subjects). The lower RMSE for the model with COM feedback compared to the default model reflects the simulated change in ankle moment in response to pelvis push and pull perturbations **(F)** that is in agreement with experimental observations **(G)**, whereas the default reflex model without COM feedback cannot capture the experimental data **(E)**. (E-G contains data of one representative subject with the two-trial average response of each unique perturbation).

We evaluated the parameter estimation results using cross-validation on novel trials at a perturbation magnitude not included in the parameter estimation. The RMSE between simulated and measured ankle moments was similar for the validation trials than for the trials used for estimation (Fig. 1. D).

### Experiment steady-state and perturbed walking with ankle exoskeleton

We implemented both controllers on a bilateral ankle-foot exoskeleton and compared them with a minimal impedance controller in twelve healthy participants during treadmill walking at 0.6 m/s. A robotic pusher was used during the experiment to apply forward and backward directed forces to a waist belt worn by the participants (Fig 2). The minimal impedance controller minimized the joint moments delivered by the exoskeleton using a disturbance observer (*27*). We used the neuromuscular controller with and without COM feedback with the identified control parameters (average across five subjects) to estimate the ankle moment based on the ankle angle measured by the exoskeleton encoders, COM velocity estimated from the trajectory of a marker on the pelvis (motion capture; only for model with COM feedback), and ground reaction forces measured using an instrumented treadmill. The exoskeleton delivered 30% of the ankle joint moment computed with the neuromuscular model.

**Fig. 2.**
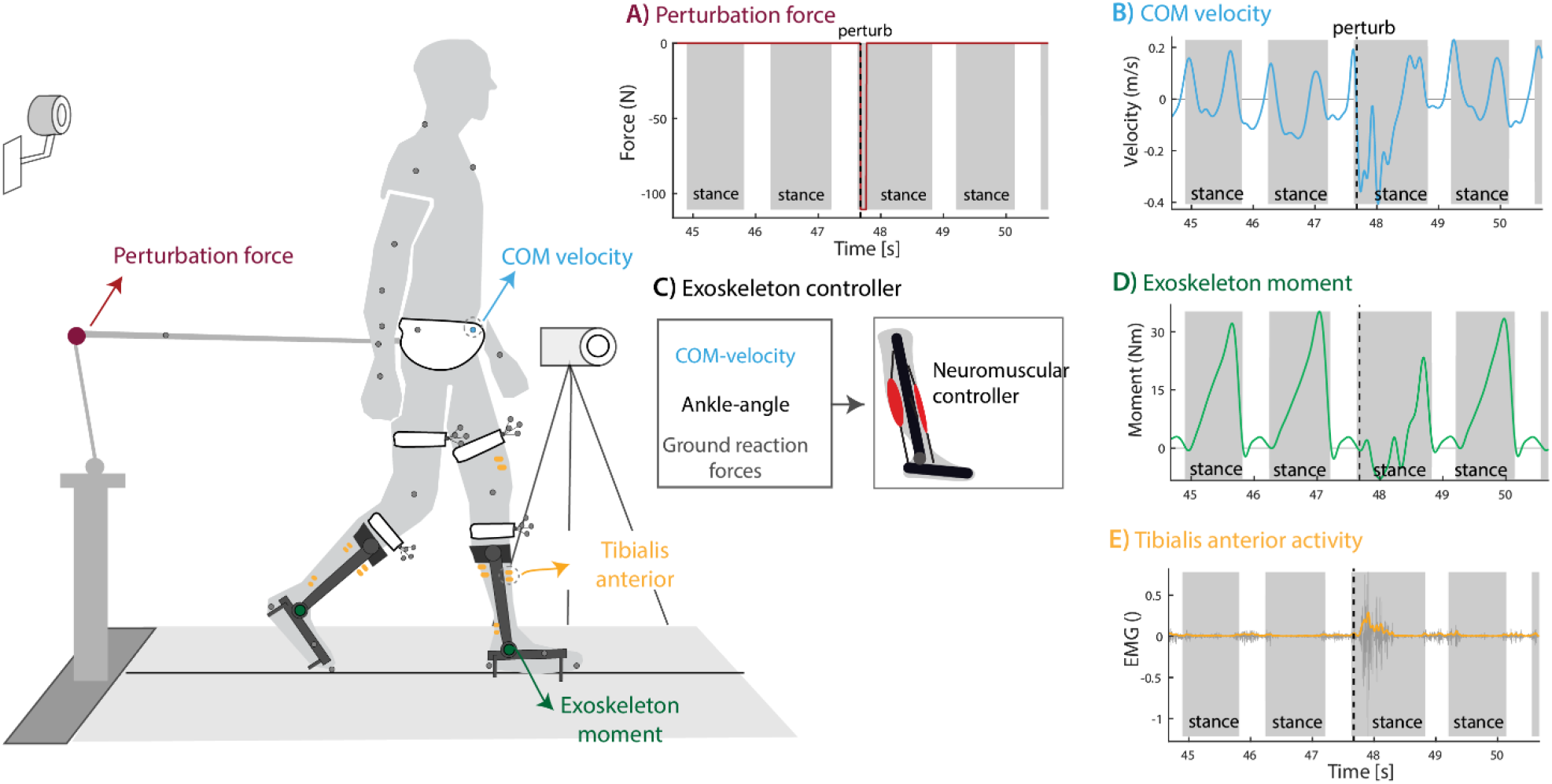
Perturbed walking with ankle-foot exoskeleton. Twelve subjects walked with a bilateral ankle-foot exoskeleton in minimal-impedance mode and controlled with the neuromechanical model with and without additional COM velocity feedback. **(A)** External forces were applied at the pelvis after right heel strike in forward (push) or backward (pull) direction to perturb human walking. **(B,C)** The neuromuscular controller uses the encoder on the ankle joint, ground reaction forces and deviation in COM velocity from the reference trajectory as input and ankle joint moment as output. **(D)** The exoskeleton delivered 30% of the ankle joint moment computed with the neuromuscular controller. **(E)** Surface Electromyography was used to quantify muscle activity to evaluate controller performance. This figure contains data of one backward directed perturbation of a typical subject walking with the neuromuscular controller with additional feedback of COM velocity.

### Muscle activity during unperturbed walking with exoskeleton

Subjects walked for 20 minutes with the default neuromuscular controller (without being perturbed) to adapt to treadmill walking with the exoskeleton. During this adaptation period, subjects changed their gait by reducing soleus activity (Fig. 3). After the adaptation period, the subjects walked for five minutes with the minimal impedance controller. On average, soleus activity was 19% lower (p = 0.001) when walking with the default neuromuscular controller than with the minimal impedance controller (Fig. 3. A-B). The decrease in gastrocnemius activity was smaller compared to the soleus (9.1, p = 0.045).

**Fig. 3.**
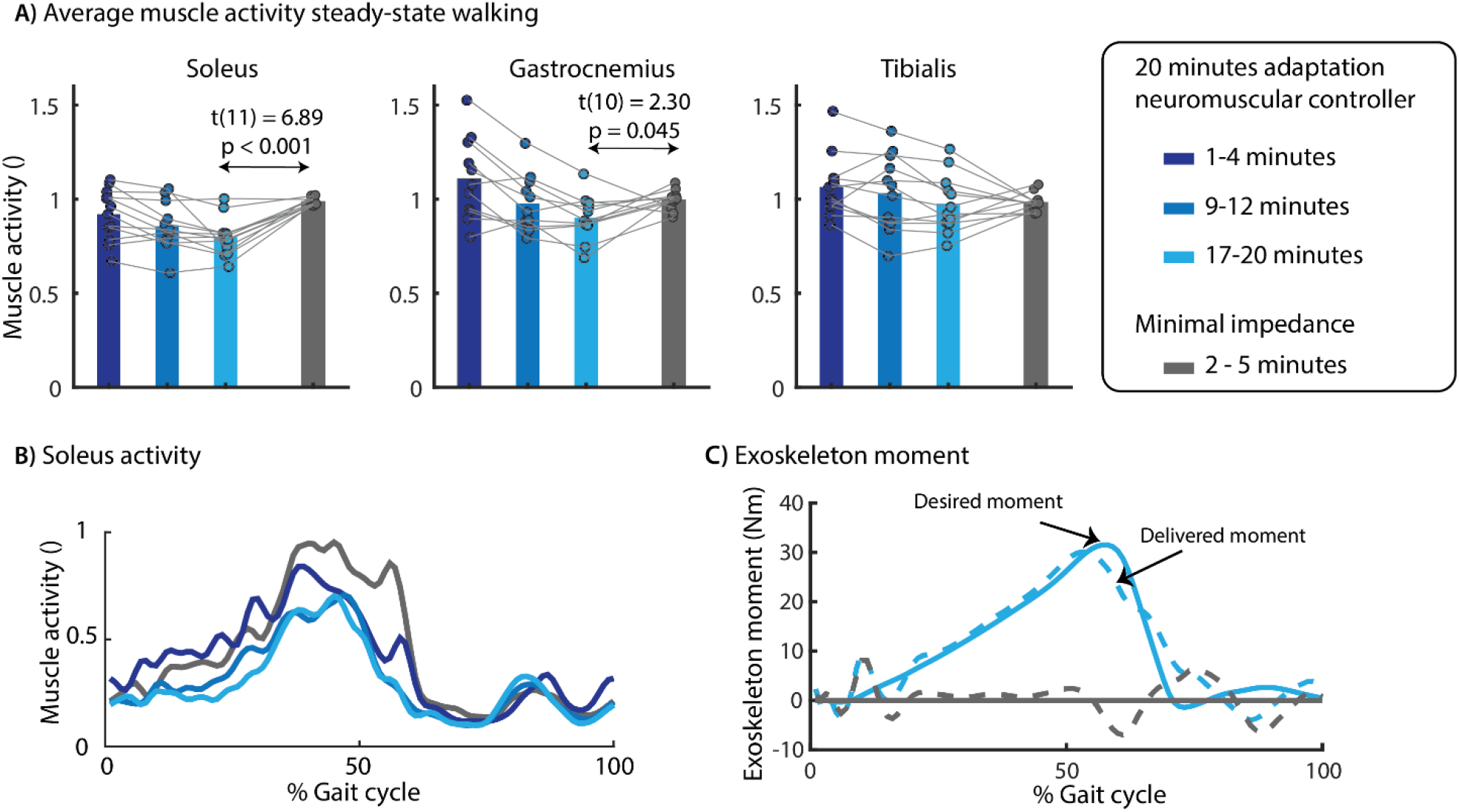
Effect of exoskeleton assistance on muscle activity in steady-state walking. We observed a gradual decrease in soleus muscle activity during the 20 minutes adaptation. Compared to minimal impedance mode (gray), average soleus activity was 19% lower at the end of the adaptation period **(A)**. The soleus activity was decreased during the full duration of the stance phase **(B)**, which is the result of the plantarflexion assistance provided by the exoskeleton **(C)**. The desired exoskeleton moment was applied with a RMSE of 2.02 Nm with mainly larger differences between desired and actual moment during the first part of the stance phase. A paired t-test was used to compare muscle activity between controllers. Panel (A) contains data of all subjects with the dots representing the median muscle activity during 3 minutes of walking and the bars the averaged data across subjects. The time series in B and C are based on data of one representative subject.

Next, subjects walked for five minutes with each of the controllers while being perturbed by forward or backward directed external forces with a magnitude of 12% of body and exoskeleton weight and a duration of 200ms applied to the pelvis. All perturbations were applied at right heel strike with randomized time intervals between perturbations. We analyzed and compared muscle activity in the last two minutes of each condition during perturbed and unperturbed gait cycles separately. During the unperturbed gait cycles, soleus activity was 20 % lower with both neuromuscular controllers than with the minimal impedance controller (Fig. S8). Hence, the neuromuscular controllers with and without COM feedback caused a similar reduction in soleus activity for unperturbed gait cycles, which is not surprising given the small variation in COM velocity during these gait cycles.

### Exoskeleton assistance in perturbed walking

Assistive joint torques computed by both neuromuscular controllers during perturbed cycles were in agreement with the simulations. The joint moments delivered by the default neuromuscular controller during perturbed gait cycle differed little from the joint moments during unperturbed gait cycle (Fig. 4.). In contrast, the joint moments delivered by the neuromuscular controller with COM velocity feedback were higher in response to pelvis pushes and lower in response to pelvis pulls than during unperturbed gait cycles (Fig. 4.). This modulation of the ankle moment with perturbation direction is similar to the human behavior observed in the experimental dataset without exoskeleton (*26*). This suggests that the subjects synergistically interact with the exoskeletons as altered responses of the subjects when walking with the exoskeletons would also alter the controller behavior, which is based on the subjects’ kinematics.

**Fig. 4.**
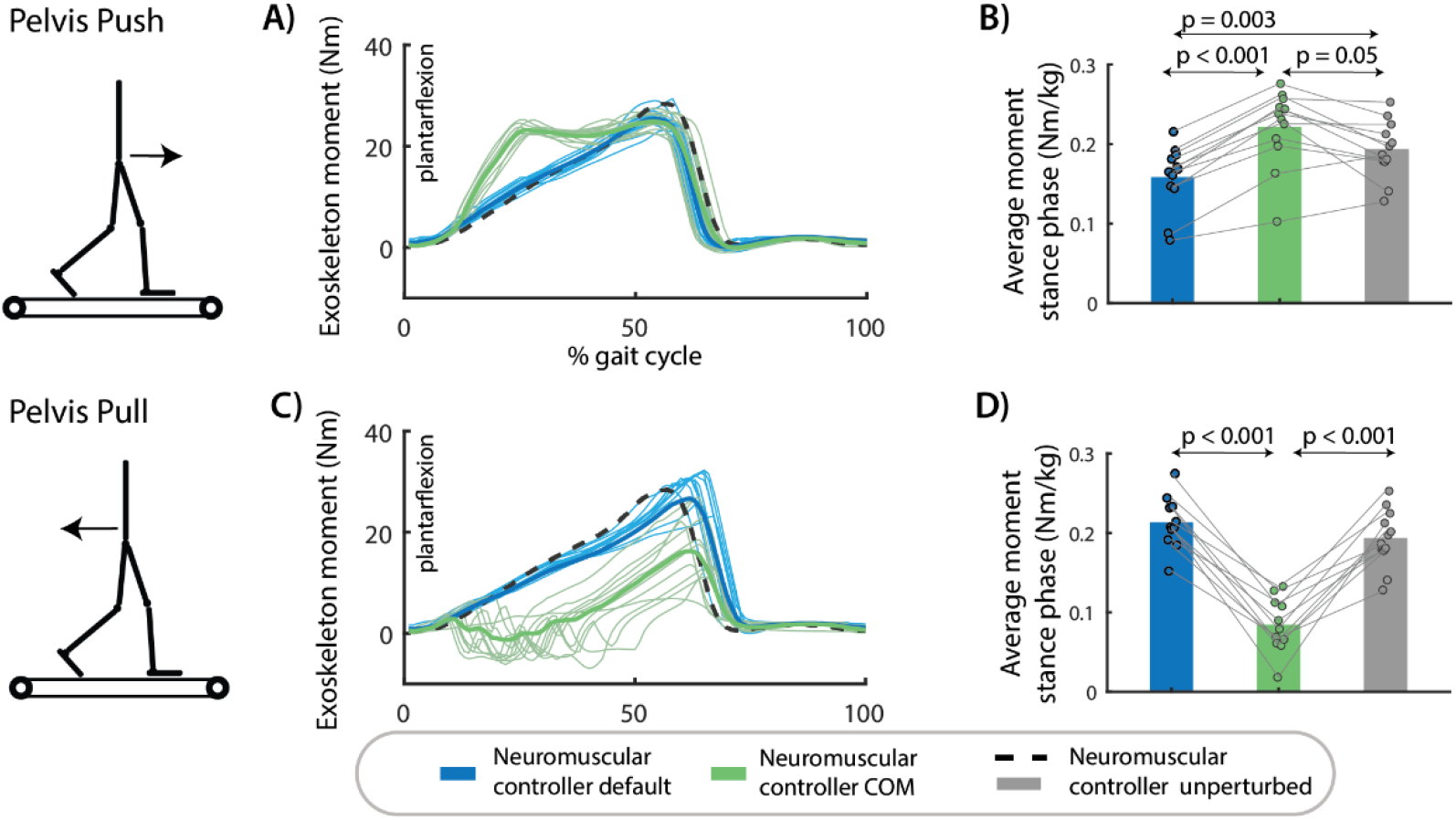
Exoskeleton assistance in perturbed walking. **(A)** The exoskeleton moment increased in response to pelvis push perturbations in the neuromuscular controller with COM feedback compared to the default neuromuscular controller and compared to steady-state walking assistance. **(B)** The type of controller had a significant influence on the average exoskeleton moment during the perturbed right stance phase after push perturbations (F(2,22) = 29.5001, p <0.001). **(C)** The assistive ankle moment decreased in response to pelvis pull perturbations in the neuromuscular controller with COM feedback compared to the default neuromuscular controller and compared to steady-state walking. **(D)** The type of controller had a significant influence on the average exoskeleton moment during the perturbed right stance phase after push perturbations (F(2,20) = 81.1381, p <0.001). A and C contain data of one representative subject. The bar plots in B and D contain data of all subjects with the dots representing the response of individual subjects and the bars the averages across subjects. A repeated measures anova with Tukey’s HSD post-hoc test was used to compare the exoskeleton moment between controllers.

### Human response to perturbation with exoskeleton assistance

The neuromuscular controller with COM feedback decreased COM displacement and muscle activity during balance recovery after perturbations compared to the default neuromuscular controller and the minimal impedance controller (Fig. 5–6).

**Fig. 5.**
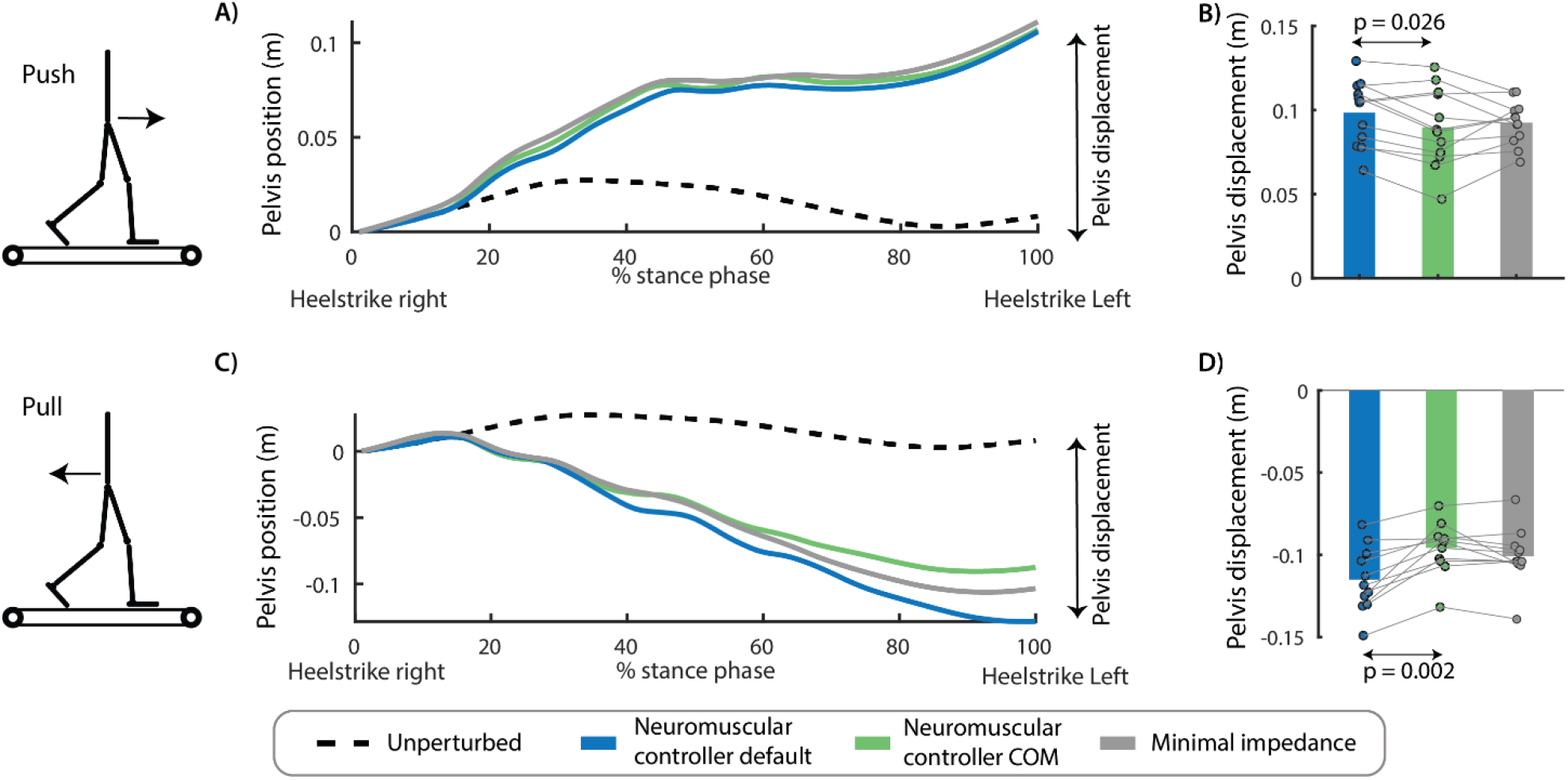
Effect of perturbation force and exoskeleton controller on pelvis displacement. The controller type influenced the movement of the pelvis after push perturbations (F(2,22) = 4.1593, p =0.04)). Forward displacement of the pelvis during the perturbed stance phase was smaller in the controller with COM feedback compared to the controller without COM feedback (p = 0.026) **(A,B)**. The controller type also influenced the backward pelvis displacement after pull perturbations (F(2,20) = 21.92, p < 0.001). Backward pelvis displacement at the first heel strike after the perturbation was smaller in the controllers with COM feedback compared to the controller without COM feedback (p = 0.002) **(C-D)**. A and C contain data of one representative subject. The bar plots in B and D contain data of all subjects with the dots representing the response of individual subjects and the bars the averages across subjects. A repeated measures anova with Tukey’s HSD post-hoc test was used to compare the pelvis displacement between controllers.

**Fig. 6.**
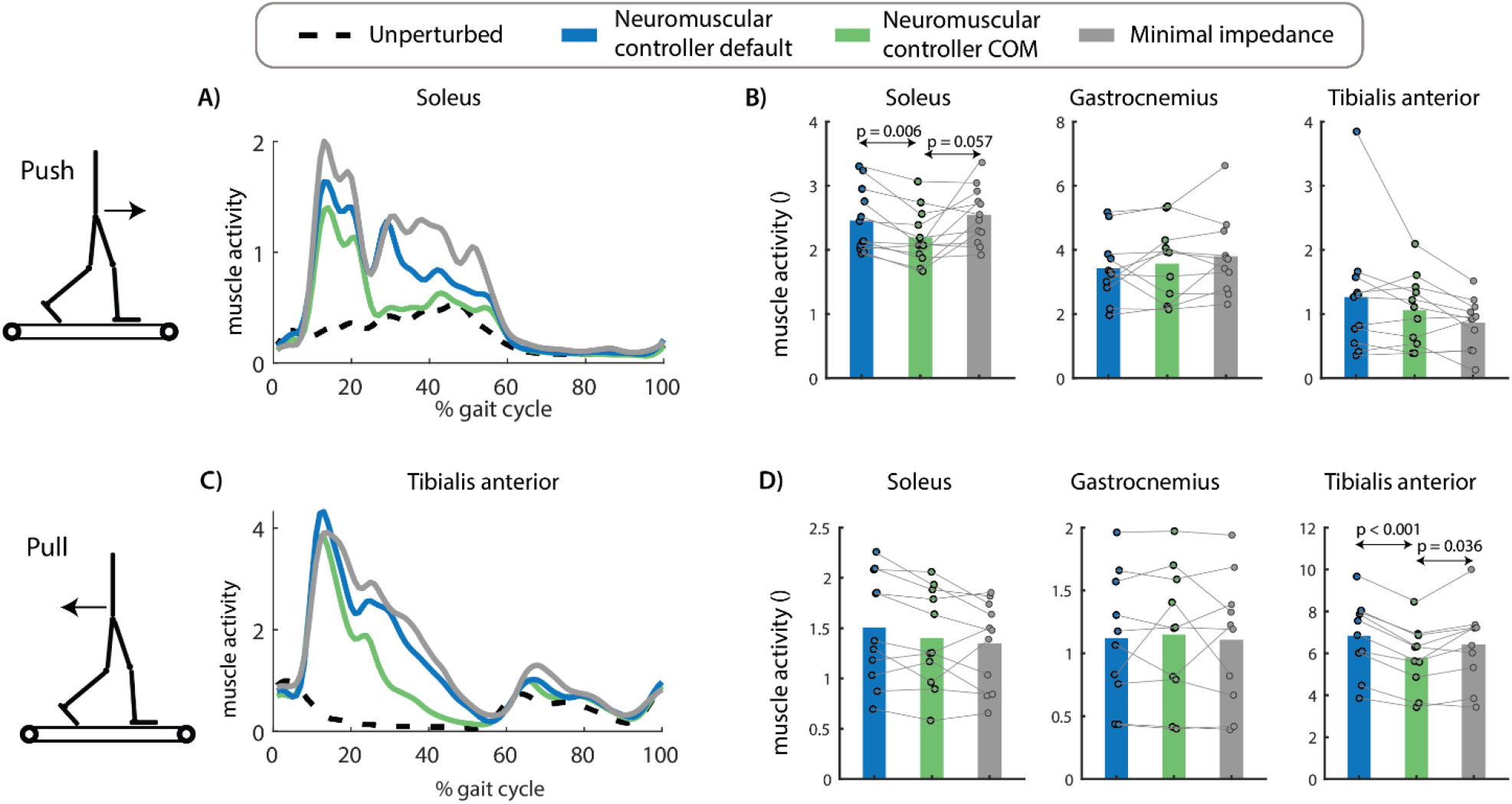
Effect of perturbation force and exoskeleton controller on muscle activity. The exoskeleton controller type influenced the increase in soleus activity after push perturbations (F(2,22) = 5.2763, p =0.013). Soleus activity in the first 500ms after perturbation onset was smaller for the neuromuscular controller with COM feedback compared to the default neuromuscular controller (p = 0.006) and compared to the minimal impedance controller (although not significant, p = 0.057) **(A-B)**. The exoskeleton controller type also influenced the increase in tibialis anterior activity after pull perturbations (F(2,18) = 15.1174, p =0.0001). Tibialis anterior activity was smaller in the neuromuscular controller with COM feedback compared to the default neuromuscular controller (p < 0.001) and compared to the minimal impedance controller (p = 0.036) **(C-D)**. A and C contain data of one representative subject. The bar plots in B and D contain data of all subjects with the dots representing the response of individual subjects and the bars the averages across subjects. A repeated measures anova with Tukey’s HSD post-hoc test was used to compare the pelvis displacement between controllers.

Push perturbations caused a forward movement of the subjects’ COM with respect to the treadmill (Fig. 5A-B) and an increase in soleus activity of the stance leg (Fig. 6A-B). The neuromuscular controller with COM feedback reduced the forward COM displacement during the stance phase with 9% compared to the default neuromuscular controller (p=0.002) and resulted in similar COM displacements than the minimal impedance controller (p = 0.828) (Fig 5 A-B). The neuromuscular controller with COM feedback reduced stance leg soleus activity during the first 500ms after the push perturbation with 10% compared to the default neuromuscular controller (p = 0.006) and with 12% compared to the minimal impedance controller (although not significant, p = 0.057) (Fig. 6 A-B).

Pull perturbations caused a backward movement of the subjects’ COM with respect to the treadmill (Fig. 5C-D) and an increase in stance leg tibialis anterior activity (Fig. 6C-D). The neuromuscular controller with COM feedback reduced the backward COM displacement during the stance phase by 18% compared to the default neuromuscular controller (p=0.002) and resulted in similar COM displacements than the minimal impedance controller (p = 0.201) (Fig 5 C-D). The neuromuscular controller with COM feedback reduced stance leg tibialis anterior activity during the first 500 ms after the pull perturbation with 16% compared to the default neuromuscular controller (p < 0.001) and with 12% compared to the minimal impedance controller (p = 0.036) (Fig. 6 C-D).

## DISCUSSION

Exoskeletons can improve the efficiency of human walking (*1–5*, *28*) but have currently limited ability to support balance control. We developed a balance supporting controller for an ankle exoskeleton that mimics the human ankle function during steady-state and perturbed walking. The controller reduced muscle activity in steady-state walking and during balance recovery after perturbations. The proposed biomimetic control strategy based on a neuromuscular model that includes muscle dynamics, local reflexes and supra-spinal balance pathways thus provides complete locomotion support including both effort and stability. This is especially important for the application of wearable robotic devices in aging and pathological populations with balance impairments.

Our control approach mimics intrinsic muscle dynamics and supra-spinal feedback control, which might both be important for its success in reducing muscle activity during balance recovery. Joint mechanical impedance is modulated during the stance phase of walking (*29*) and has important contributions to stabilize human walking (*30*). Joint impedance originates both from muscle mechanical properties and muscle control. In our model, mechanical impedance originates from the force-length-velocity relationship in the virtual Hill-type muscle (*31*). Mechanical impedance has been shown to be important to reduce the need for active control through delayed feedback (*30*, *32*, *33*). However, mechanical impedance and local reflexes alone cannot explain observed changes in ankle torque after perturbations (Fig. 1, default neuromuscular model). The comparison of the controller with and without COM velocity feedback demonstrates that supra-spinal feedback is important for supporting balance control during walking with wearable robotic devices. The importance of task-level feedback is in line with previous research on the use of an ankle exoskeleton to support perturbed standing balance, where reductions in COM movement and muscle activity were larger with a controller that used feedback from body sway and sway velocity than local feedback from ankle angle and angular velocity (*34*). We chose to also mimic human feedback delays. In general, controller performance decreases with increasing delays. Therefore, we might be able to further reduce muscle activity and COM displacement by reducing feedback delays. It should be investigated how such augmented balance control influences performance and interaction with the human.

Our results suggest that human sensorimotor processing was unaltered by exoskeleton support. Similar as in perturbed walking without an exoskeleton (*21*), we found that humans use COM feedback in response to perturbations. Specifically, variability in the muscle response to perturbations can be explained by variability in COM movement (Fig. S9-10). We therefore infer that the reduction in COM excursion and velocity with the novel exoskeleton controller induced reductions in muscle activity in the absence of large changes in human balance control.

It remains to be tested whether the proposed controller can support walking balance in patients with altered sensorimotor transformation underlying balance control. It has been documented that sensorimotor processing underlying standing balance control is altered in older adults and patients with Parkinson’s disease (*35*). In contrast to healthy adults, they also recruit antagonistic muscles in response to center of mass perturbations during standing (*35*). It is largely unknown how sensorimotor processing underlying walking balance control is affected in persons with neurological disorders. It is therefore unclear whether patients with balance control deficits can also exploit the provided balance support. Whereas it seems unlikely that adapting the controller to mimic the balance control strategy of mobility impaired individuals would improve balance, it is unclear whether the control strategy inspired by healthy adults should be adapted for mobility impaired individuals.

Extending the proposed balance controller to support balance across walking conditions is expected to be straightforward given that the underlying COM feedback strategy explains human balance control across conditions. We previously demonstrated that COM feedback can explain corrective ankle moments across perturbation types (pelvis perturbations and support-surface translations) and gait speeds (*21*). However, COM velocity feedback gains change with gait speed and throughout the stance phase. Faster walking is mechanically more stable and relies more on adjustment of foot placement to control balance due to the higher step frequency (*24*). Hence, balance control during faster walking relies less on the modulation in the ankle moment. Implementing a controller with feedback gains that are modulated with gait speed and gait phase is feasible given that both can be estimated from wearable sensors (e.g. inertial measurement unit (*36*). Adaptability of the balance correcting feedback pathways would complement the adaptability of the local feedback pathways. It has been shown that the default controller is able to reproduce steady state walking at various speeds in simulations (*17*) and results in speed-adaptive behavior when used to control a transtibial prosthesis (*37*).

Reductions in muscle activity do not imply reductions in metabolic energy consumption. It is hard to relate the observed 19% reduction in soleus activity during steady-state walking to a reduction in metabolic power. Metabolic power does not only depend on muscle activity but also on the operating conditions of the muscle. Previous studies have demonstrated that an external force provided by the exoskeleton in parallel with the compliant muscle-tendon unit can alter the operating length and velocity of the muscle and therefore undermine the energy efficiency (*38*, *39*). We expect that our controller can be further optimized to reduce metabolic cost. Previous research has demonstrated that the timing of assistance is important to reduce the metabolic cost (*3*). Exoskeletons that were successful in reducing metabolic cost mainly provided assistance in late stance, when the muscle fibers are performing metabolically costly positive work. However, simultaneously reducing metabolic power and improving stability might require dedicated strategies.

Our observation that muscle activity is reduced with the default neuromuscular controller is in contrast with previous work. Shafer et al. found an increase in soleus activity during early stance and swing and an increase in metabolic power with a similar controller (*16*). Multiple differences between study protocols may explain the discrepancy between both studies. First, Shafer et al. tested the controller at a walking speed of 1.25 m/s in (*16*) whereas our participants walked at 0.6 m/s. Second, we slightly modified the local feedback controller. We implemented a gradual change in feedback gains between stance and swing phases and identified control parameters based on experimental data whereas Shafer et al. (*16*) only did a sweep of two control parameters. A more gradual change in control parameters allowed us to better capture the biological torque. Without these adaptations, we overestimated the ankle torque in early stance, which might explain the increase in soleus activity during early stance in the experiments of Shafer et al. (*38*) (Fig. S3).

Implementing the proposed controller in daily life requires wearable alternatives for the lab-based sensors, but we believe that these sensor are readily available. Laboratory-based sensors to measure the ground reaction force and COM velocity were used as input in the controller. We believe that these sensors can be easily replaced by foot switches to detect contact with the ground and an inertial measurement unit to estimate COM velocity (*40*). We tested our controller on a wearable exoskeleton that was originally designed to support individuals with complete spinal cord injuries (*41*). As a result, the exoskeleton was over-dimensioned for our study, explaining the relatively high weight (5 kg on each ankle-foot). For this reason, we did compare our controller’s performance to a minimal impedance controller rather than to walking without an exoskeleton. However, our controller is not device-specific and therefore applicable to other hardware designs. Given sufficiently light hardware, we expect our controller to reduce muscle activity with respect to walking without an exoskeleton but this remains to be demonstrated. Our current results can thus best be seen as a proof of principle for a biomimetic control design for balance support.

## MATERIALS AND METHODS

### Data for controller parameter identification

We first identified control parameters of a neuromechanical model based on an existing motion-capture dataset with pull and push perturbations applied at the pelvis at toe-off of the contralateral leg during walking at 0.62 m/s (details in (*26*)). In summary, steady-state walking was perturbed by means of an external force applied at the pelvis in the walking direction (pelvis push) or in the opposite direction (pelvis pull) with four different magnitudes (perturbation pulse of 150ms ranging between −0.16 to 0.16% body weight). Joint kinematics and kinetics were computed using a scaled generic musculoskeletal model with 23 degrees of freedom (gait 2392) in OpenSim (*42*). This model was scaled to the anthropometry and mass of the subject based on the marker positions and ground reaction forces in a static trials. Joint kinetics were computed based on the equations of motion of the model with OpenSim’s inverse dynamics tool.

### Neuromechanical model

We modeled the ankle moment as the sum of the moment of a mono-articular plantarflexor muscle (i.e. mimicking the soleus) and a dorsiflexor muscle (i.e. mimicking the tibialis anterior). We approximated the relation between the ankle angle, and muscle-tendon length and moment arms from the gait2392 using polynomial functions (*42*, *43*). The Hill-type muscle dynamics were implemented as in (*44*) with activation dynamics, an elastic tendon, a parallel passive element and a contractile element with a force-length and force-velocity relationship. The maximal isometric force of the plantarflexor muscle was adjusted to represent the combined force generating capacity of the gastrocnemius and soleus (see appendix table S1 for details).

Both muscles were driven by gait-phase dependent reflexes according to the model proposed by Geyer et al. (*12*). The soleus reflex consists of delayed (*τ*_*m*_= 30 ms) positive force feedback during the stance phase with gain (*G*_*sol*_) and baseline activity (*e*_*sol*,0_) (eq. 1). The tibialis anterior reflex consists of baseline activity (*e*_*ta*,0_), length feedback with feedback gain *G*_*ta*_, and inhibition proportional to soleus force *G*_*sol*,*ta*_during the stance phase (eq. 2).

To test the hypothesis that supra-spinal feedback is needed to model the change in ankle moment after perturbation, we included an additional reflex with delayed (*τ*_*m*_= 60 ms) feedback of deviations in COM velocity in the walking direction 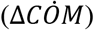 during the stance phase with feedback gain (*K*_*sol*_) for the soleus and (*K*_*ta*_) for the tibialis anterior. Deviations in COM velocity were computed as the difference between COM velocities expressed as a percentage of the gait cycle after the perturbation and during steady-state walking.

Finally, we found that the fit between simulated and experimental ankle moments could be improved when implementing a gradual transition between the stance and swing phase feedback gains. This gradual transition was implemented based on the vertical ground reaction force (*F*_*z*_) (eq. 3).

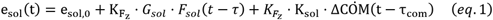

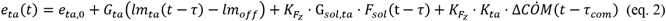

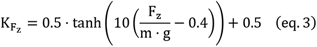

with *m* the total body mass of the subject, *g (=9.81 m/s^2)* the standard acceleration due to gravity and 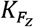 the scale factor that equals 1 during the stance phase and is zero during the swing phase. The appendix contains more details on the neuromechanical modeling.

### Controller parameter identification

We estimated control parameters that can describe the ankle moment during both steady-state walking, and walking with pelvis push and pull perturbations by minimizing the differences between the experimental (inverse dynamics) and simulated ankle moment across trials.

We estimated the eight control parameters (*e*_*sol*,0_, *G*_*sol*_, *K*_*sol*_, *e*_*ta*,0_, *G*_*ta*_, *lm*_*off*_, G_*sol*,*ta*_, *K*_*tib*_) by simultaneously minimizing the tracking error over eight steady-state gait cycles and 16 perturbation trials (two directions, four magnitudes and two repetitions of each perturbation). This ensures that we find control parameters that can describe both normal and perturbed walking. Note that common feedback gains were estimated for push and pull perturbations as direction-dependent gains resulted in only 0.33 Nm and 1.26 Nm decrease in RMSE for respectively pelvis push and pull perturbations (*Fig. S4*). Also note that Hill-type muscle properties and model time-delays were not optimized to avoid overfitting.

All optimization problems were formulated using a shooting approach in Casadi (*45*) and solved using ipopt (*46*). The forward simulation of muscle dynamics was implemented in matlab using a forward euler integration scheme with a constant step size of 0.001s. The measured inverse dynamic moment and ankle moment computed with the neuromuscular model was scaled based on the mass and height of an average subject (mass = 70 kg, height = 1.75m) to facilitate comparison of the reflex parameters between subjects.

The interdependence of the control parameters was evaluated using a method proposed by (*47*). We found that two reflex parameters (baseline soleus activity and soleus force feedback gain) were highly correlated in this dataset. Therefore, we decided to keep the baseline soleus activity constant (*e*_*sol*,0_ = 0.027) during the final parameter estimation process (Fig. S5).

Cross-validation of the estimated feedback gains was performed by predicting a novel perturbation magnitude (that was not included in the parameter estimation). We predicted ankle-joint moments for perturbations of 12% body weight with feedback gains estimated on perturbations magnitudes of 4, 8 and 16% body weight.

We evaluated the fit between measured (inverse dynamics) and simulated ankle moments by the root mean square error between both. We evaluated the RMSE separately for perturbed and steady-state gait cycles (Fig. 1). The similarity between simulated and measured (with electromyography) muscle activity was evaluated qualitatively (Fig. S2).

### Participants

Twelve healthy participants (age: 29 +/− 5 years, body mass: 69.28 +/− 8.67 kg, height: 1.73 +/− 0.08 m; mean +/− SD) took part in the experiments with the ankle-foot exoskeleton. We only included participants without a history of musculoskeletal or neurological disorders. The participants did not receive information about the different exoskeleton controllers that were tested. Subjects wore a safety harness with a fall protection system for the entire duration of the experiment. All participants provided written informed consent. The experimental protocol was approved by the local ethical committee of the University of Twente (reference number 2020.30).

### Bilateral ankle-foot exoskeleton

The left and right ankle modules of the symbitron exoskeleton (*41*) were used to assist plantar-dorsiflexion during steady-state and perturbed walking. Each ankle module weighs 5 kg. The series elastic actuator can deliver a controlled peak moment of 100 Nm and has a maximum output speed of 5 rad/s. Motor position and joint position are measured via encoders. A control computer executes the controller in TwinCat in real-time with a sampling frequency of 1 kHz.

We tested three controllers. The first controller is a minimal impedance controller described elsewhere (*25*). This controller relies on a disturbance observer to lower the apparent impedance. The second and third controller delivered assistance in combination with the minimal impedance controller. The desired exoskeleton assistance was set to 30% of the estimated subject’s biological moment computed with the default neuromuscular model and the neuromuscular model with COM feedback. The value of 30% was chosen to have a similar peak exoskeleton moment as in an experiment that optimized exoskeleton assistance to reduce the metabolic energy consumption during walking (*2*). The inputs to the neuromuscular model were the ankle angle measured by the exoskeleton encoders, the ground reaction forces measured by the instrumented split-belt treadmill, and the COM velocity estimated based on the trajectory of a marker on the pelvis. Unfiltered vertical ground reaction forces, measured with the split-belt treadmill, were used in real-time to modulate the phase-dependent reflex gains in the neuromechanical model. An optical motion capture system (Qualisys, Göteborg, Sweden) was used to estimate the COM kinematics in real-time. A single marker on the pelvis brace (Fig. 2) was used to approximate the motion of the COM in real-time. The marker position was differentiated with respect to time and band-pass filtered in real-time (IIR filter, 5–50 Hz, 0.140 dB, Simulink 2018). Deviations in COM kinematics from steady-state trajectories were computed using a simple single learner (i.e. look-up table). The average COM trajectory as a function of the gait cycle was continuously updated as it was computed from the previous six unperturbed gait cycles prior to the perturbation. This average trajectory was used to predict the current COM velocity in the anterior-posterior direction using interpolation. The deviation in COM velocity was defined as the difference between the measured COM velocity (from optical motion capture) and the COM velocity from the signal learner.

### Protocol perturbed walking with exoskeleton

The participants walked on a dual-belt treadmill (Y-mill, Motek Medical, Amsterdam, The Netherlands) at a constant speed of 0.6 m/s. We selected this slow walking speed because; (a) this is similar to the walking speed in the perturbation experiments that we used for parameter estimation (*26*), (b) large changes in ankle moment are observed in response to pull and push perturbations at this speed (*21*, *26*), and (c) faster walking speeds are more challenging for the subjects with the heavy exoskeleton. The experiment started with a 20 minute walking trial without perturbations with the default neuromuscular controller to adapt to the treadmill, exoskeleton geometry, added mass of the exoskeleton and the exoskeleton assistance. Subsequently the participants walked five minutes with the exoskeleton in minimal impedance mode. After a 10-minute rest, the participants walked four trials of five minutes each with perturbations, where four exoskeleton controllers were tested: minimal impedance controller, default neuromuscular controller, neuromuscular controller with COM velocity feedback, and a fourth controller that was not included in this study. The order of the four trials was randomized for each participant. Perturbations were applied in anterior and posterior direction using a pusher device (Moog, Nieuw-Vennep, Netherlands) attached to the subjects’ pelvis by a soft brace (*26*). The pelvis was chosen as the point of application of the external perturbation, as it approximately coincides with the location of the whole-body COM.

Perturbations were applied at right heel contact (when the right leg vertical ground reaction force exceeded 50N). The perturbation consisted of a square force pulse of 0.2s and a magnitude of 12% of combined body and exoskeleton weight and was semi-randomly applied in anterior and posterior direction. The time between perturbations was semi-randomized to prevent anticipation and varied between 8s and 16s, resulting in a total of 22 perturbations (11 push, 11 pull) during the 5-minute trials.

### Data acquisition

Kinematic data of bony landmarks on the feet, ankles, knee, pelvis and torso and cluster markers on the tibia and femur were recorded at 128 Hz using an optical motion capture system with 8 Oqus cameras (Qualisys, Göteborg, Sweden). Note that we only used kinematic data of markers on the feet and pelvis in the data analysis. Ground reaction forces were collected on a split-belt treadmill with a sampling frequency of 2048Hz (Y-mill, Motek Medical, Amsterdam, The Netherlands). Muscle activity of the left and right soleus (Sol), gastrocnemius lateralis (Gas) and tibialis anterior (Tib) was measured using surface electromyography (Bagnoli, Delsys, Natcik, MA, USA), sampled at 2048Hz. Data related to the exoskeleton controller (encoder values, desired moment, applied moment, …) and the pusher (perturbation onset) were logged at 1000 Hz through the exoskeleton computer. All data were synchronized using the ground reaction forces, whose analog signals were logged by both the exoskeleton and Qualisys computers.

### Data processing

Data were processed in Matlab 2021 (Mathworks, Natick, MA, USA). EMG data were filtered with a second order IIR notch filter to remove the electric hum and a 2nd order, zero-lag, Butterworth bandpass filter with cut-off frequencies of 20 and 400 Hz. The filtered signals were rectified and a linear envelope was created with a 2nd order, zero-lag, Butterworth low-pass filter with a cut-off frequency of 20 Hz. The filtered EMG data was normalized based on the average muscle activity in the five minutes steady-state walking in minimal impedance mode.

### Outcomes

The moment delivered by the exoskeleton was used to assess the assistance provided by the exoskeleton during steady-state and perturbed walking. The exoskeleton moment was expressed as a percentage of the gait cycle to quantify the assistance during steady-state walking and the modulation of the assistive moment in response to the perturbations.

We evaluated if the assistance provided by the exoskeleton reduced muscle activity during steady-state walking. We compared the average muscle activity during the last two minutes of the steady-state walking sessions with the neuromuscular controller and the minimal impedance controller. In addition, we also compared the average muscle activity of the unperturbed gait cycles during the last two minutes of the perturbation session between controllers to verify that further adaptation of subjects to the controller during the perturbation experiment or anticipation to the perturbations did not affect performance.

We evaluated whether the controller influenced the muscle activity and COM movement in response to the perturbation. We compared the average muscle activity during the first 500ms after perturbation onset to evaluate muscle activity during balance recovery in each controller. The time window of 500ms after perturbation was selected as this includes the main changes in muscle activity (Fig 6 A-C) and this time window was also used in a similar study (*10*). The movement on the treadmill was computed as the displacement of the pelvis marker from perturbation onset (right heel strike) until the subsequent left heel strike. For both the muscle response to the perturbation and COM displacement, we computed the median of each outcome of the 11 repetitions of push and pull perturbations for each subject.

### Statistical analysis

A two-sided paired t-test with an alpha = 0.05 was used to evaluate if the exoskeleton reduced muscle activity during steady-state walking as the data was normally distributed (Mauchly’s sphericity test). We used a repeated measures anova to evaluate if the type of controller influenced the muscle activity and COM movement in response to the perturbation. A Greenhouse-Geisser correction was applied in case of a lack of sphericity in the data, indicated by Mauchly’s test for sphericity. When anova test was significant, Tukey’s Honestly Significant Difference was employed as a post-hoc test to compare the three controllers.

## Acknowledgements

The authors would like to thank Michiel Ligtenberg, Robert Roos and Lars D’hondt for the technical support with hardware and software.

## Data and code

The data of the perturbation experiment with the ankle-foot exoskeleton and the software for parameter identification, control of the exoskeleton and analysis of the experimental data will be shared after peer-review.

