## Supplementary information for "Assisting walking balance using a bio-inspired exoskeleton controller"

### Supplemental material for: Assisting walking balance using a bio-inspired exoskeleton controller

### 1 Neuromechanical model

#### 1.1 Muscle dynamics

A standard Hill-type model with a series elastic element (tendon) and contractile element with active and passive properties (muscle fibers) was used to model the muscle-tendon dynamics of the soleus and tibialis anterior. We used the mathematical model of muscle dynamics as described in (44). In short, the length of the muscle tendon unit ($l_{MTU}$) was a function of the modeled musculoskeletal geometry and ankle angle ($\phi_{ankle}$) as modeled in the Opensim *gait23dof92musc* model.

$$l_{MTU} = f(\phi_{ankle}) \tag{1}$$

$$l_T = l_{MTU} - l_m \cdot cos(\alpha) \tag{2}$$

with $\alpha$ the pennation angle of the muscle. The pennation angle was computed under the assumption of a constant width $l_w$ of the muscle fiber that is equal to $l_w = l_{m,opt} \cdot sin(\alpha_0)$. With $\alpha_0$ the pennation angle at optimal fiber length.

| | soleus | tibialis |
|---|---|---|
| Maximal isometric force ($F_{iso}$) | 7000 N | 1500 N |
| Optimal fiber length ($l_{m,opt}$) | 0.06 m | 0.0972 m |
| Tendon slack length ($l_{Ts}$) | 0.249 m | 0.2011 m |
| Pennation angle at $l_{m,opt}$ ($\alpha$) | 0.436 rad | 0.0873 rad |
| Maximal fiber velocity ($\dot{l}_{m,max}$) | 0.6 m/s | 0.97 m/s |
| Scale factor tendon F/L ($k_T$) | 20 | 35 |

Table S 1: Muscle properties of the soleus and tibialis anterior muscle

The force in the muscle tendon unit ($F_T$) is a nonlinear function of the tendon strain.

$$F_T(\tilde{l}_T) = F_{iso}(\frac{1}{5} \cdot e^{k_T(\tilde{l}_T - 0.0995)} - c_1) \tag{3}$$

with $\tilde{l}_T$ the length of the tendon divided by the tendon slack length $l_{Ts}$, and $F_{iso}$ is the maximal isometric force of the muscle. The parameters $k_T$ and $c_1$ where chosen to represent the force-length relation observed in the Achilles tendon and in the tendon of the tibialis anterior with $k_T = 20$ and $c_1 = -0.2328$ for the soleus and $k_T = 35$ and $c_1 = -0.25$ for the tibialis anterior. Contractile element velocity was calculated from muscle force-length and force-velocity and activation of the muscles (equation 4)as described in (16). The parameters of the plantarflexors and tibialis anterior were based on the OpenSim *gait23dof92musc* model that was slightly adjusted to combine the force generating capacity of the soleus and gastrocnemius (table 1). A forward euler integration scheme was used to calculate the length of the contractile element in the next time step *(i+1)*.

$$\dot{l}_m = f(l_m, a) \tag{4}$$

$$l_{m,i+1} = l_{m,i} + \dot{l}_{m,i} \cdot \Delta t \tag{5}$$

with $\Delta t = 0.001\ s$.

#### 1.2 Neural control policy

The muscle model was driven by baseline muscle excitation and local feedback of muscle states in accordance with an existing model (12) and feedback of center of mass velocity (equation 6 and 7). Control parameters were optimized to track inverse dynamic ankle joint moments, which resulted in a set of control parameters used to control a bilateral ankle foot exoskeleton (table 2). We implemented a gradual transition between stance and swing phase feedback gains, based on the vertical ground reaction forces (equation 8), to improve the fit between inverse dynamic and simulated joint moments.

$$e_{sol}(t) = e_{sol,0} + K_{F_z} \cdot G_{sol} \cdot F_{sol}(t - \tau_m)) + K_{F_z} \cdot K_{sol} \cdot \Delta \dot{COM}(t - \tau_{com}) \tag{6}$$

$$e_{ta}(t) = e_{ta,0} + G_{ta}(lm_{ta}(t - \tau_m) - l_{m,off})) + K_{F_z} \cdot G_{sol,ta} \cdot F_{sol} + K_{F_z} \cdot K_{tib} \cdot \Delta \dot{COM}(t - \tau_{com}) \tag{7}$$

$$K_{F_z} = 0.5 \cdot tanh(10(\frac{F_z}{m \cdot g} - 0.4)) + 0.5 \tag{8}$$

| Soleus | | Tibialis anterior | |
|---|---|---|---|
| $e_{sol,0}$ | 0.027 | $e_{ta,0}$ | 0.02 |
| $G_{sol}$ | 1.28 | $G_{ta}$ | 1.49 |
| $K_{sol}$ | 0.32 | $l_{m,off}$ | 0.9 |
| | | $G_{sol,ta}$ | 0.45 |
| | | $K_{ta}$ | -1.65 |

Table S 2: control parameters for the soleus and gastrocnemius muscle

with $K_{F_z}$ the scale factor between 0 and 1 causes a smooth transition between stance and swing reflexes, $F_z$ the vertical component of the ground reaction force, $m$ the subject mass, $g = 9.81\ m/s^2$ the gravitational acceleration.

### 2 Supplementary information parameter identification

#### 2.1 Overview optimization problem

We estimated the control parameters of each model to track the inverse dynamic joint moment in eight steady-state gait cycles and 16 perturbation trials (two perturbation directions, four perturbation magnitudes and two repetitions of each perturbation) in one optimization problem per subject. The difference between the inverse dynamic joint moment and simulated moment is visualized in Fig. S1 for the neuromusclar controller with and without COM velocity feedback.

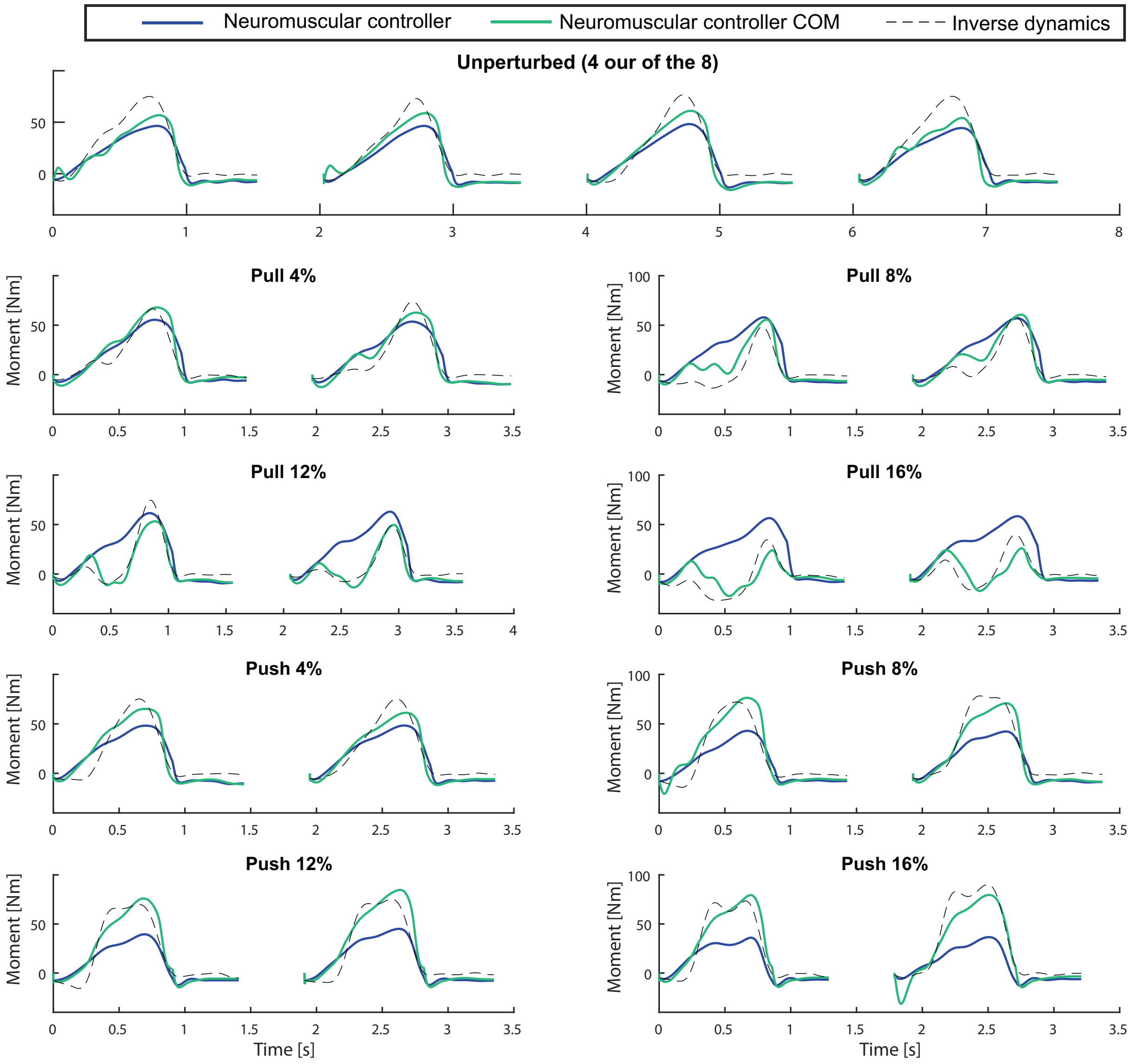

Figure S 1: **Overview of the parameter estimation**. Inverse dynamic and simulated ankle joint moments for all gait cycles included in the optimization problem in one representative subject.

#### 2.2 Predicted muscle activity

Although we did not optimize the fit between measured and simulated muscle activity, the model with additional COM feedback also predicts an increase in calf muscle activity in response to push perturbations and an increase in tibialis anterior activity in response to pull perturbations (Fig. S 2). The initial muscle response to the perturbations (i.e. first 200 ms) was however larger in the experiment compared to the simulations, which indicates that additional feedback of center of mass accelerations (as proposed by (22)) might improve the model further.

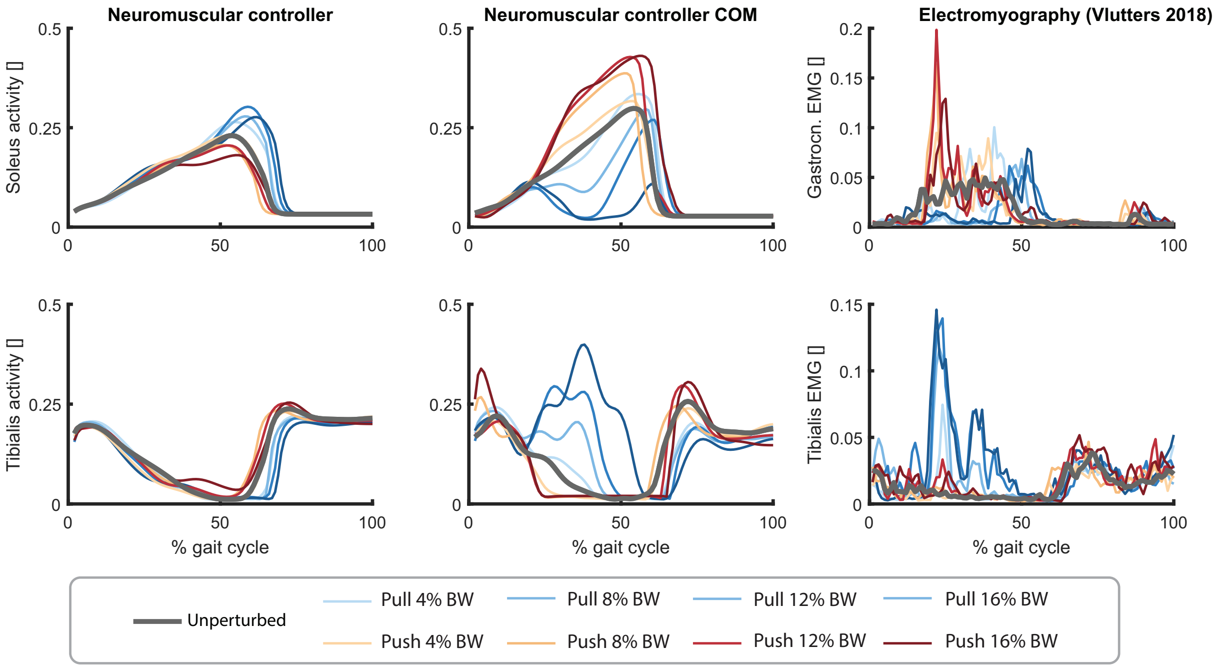

Figure S 2: **Measured and simulated muscle activity in parameter estimation.** The neuromuscular model with additional feedback of COM velocity can simulate the modulation of soleus and tibialis anterior muscle activity after pelvis push and pull perturbations. This figure contains simulated muscle activity of the soleus (top row) and tibialis anterior (bottom row) in response to pelvis pull (blue) and pelvis push (red) perturbations of different magnitudes (shades of the color) in one typical subject. We compared the default neuromuscular controller (left column) with neuromuscular controller with additional feedback of COM velocity (middle column) and electromyography data (right column). The model with COM feedback predicts the increase in calf muscle activity in response to push perturbation and increase in tibialis anterior activity in response to pull perturbations, which is not simulated with the model without COM feedback. The model with COM feedback also predicts the decrease in Soleus activity in response to pull perturbations. Note that there is no electromyography data on soleus muscle activity and was therefore compared to electromyography data of the gastrocnemius.

#### 2.3 Smooth transition between stance and swing control

We found that the fit with experimental data could be improved when implementing a gradual change between the stance and swing phase feedback gains. The original neuromuscular controller proposed by (12) has a discrete transition between the stance and swing phases with $G_{sol} = 0$ and $G_{ta} = 0$ during the swing phase. We observed that a gradual transition between stance and swing reflex gains reduced the RMSE in ankle torque 2.23 Nm (Fig. S3).

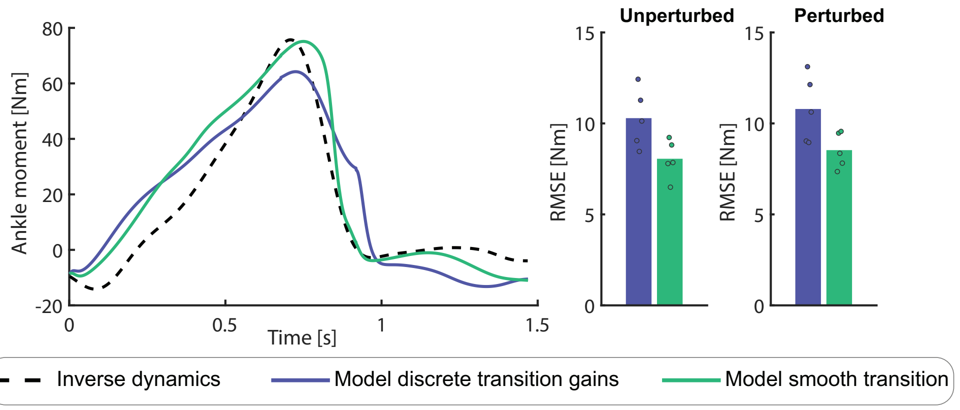

Figure S 3: **Smooth transition between stance and swing control.** The neuromuscular model with a smooth transition between stance and swing reflex gain (green) can track the inverse dynamic joint moment more closely compared to the default model with a discrete change in reflex gains (blue). The left figure shows a the ankle moment for a typical subject and unperturbed gait cycle. The right bar plots show the RMSE for all subjects in steady-state and perturbed walking.

#### 2.4 Direction dependent reflex gains

We investigated whether separate reflex gains ($K_{sol}$ and $K_{tib}$) for push and pull perturbations improves the fit between measured and simulated ankle joint moments. We found that separate reflex gains decreased the RMSE between measured and simulated joint moment with only 1.26 Nm for pelvis pull and with 0.33 Nm for pelvis push perturbations (Fig S4). Hence, we decided to use the simplest model with one set of gains independent of perturbation direction.

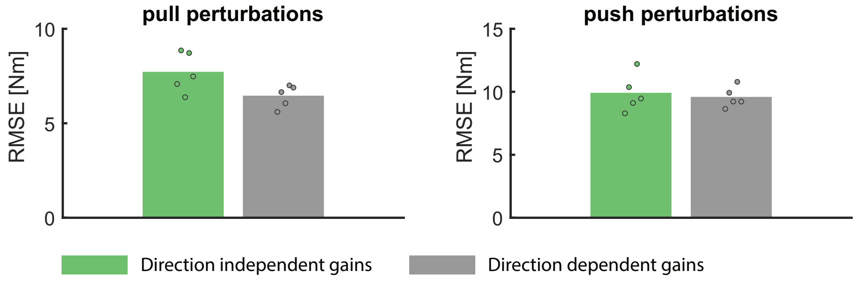

Figure S 4: **Direction dependent reflex gains** decreased the RMSE between simulated and inverse dynamic ankle joint moment only slightly in response to pull and push perturbations. Hence a model with one set of gains for forward and backward deviations in COM velocity (i.e. push and pull perturbations) was implemented

#### 2.5 Interdependence of control parameters

The covariance matrix of the control parameters (P) was derived to determine the interdependence of the control parameters of the neuromuscular model (equation 9).

$$P = \frac{1}{N} \cdot (J^T \cdot J)^{-1} e \cdot e^T \tag{9}$$

With N the number of time steps, J the Jacobian of the cost to the parameters and e is a vector with the difference between measured and simulated joint moments and each time step. The Jacobian is a N x np matrix, with np is the number of estimated parameters (i.e. np = 6 for the default neuromechanical model and np = 8 for the neuromechanical model with COM feedback). The interdependence of the parameters was evaluated by comparing the auto-covariance (diagonal terms of P) to the cross-covariance (off-diagonal terms of P).If the auto-covariance was higher than all cross-covariances, the corresponding parameter was estimated/assumed independently and its estimated value was assumed to be reliable. Similar as in (45), we normalized the covariance matrix such that all diagonal terms (auto-covariance) equal one ($P_{i,j}^{plot} = |\frac{P_{i,j}}{\sqrt{P_{i,i} \cdot P_{j,j}}}|$) to facilitate graphical interpretation of the results (Fig S 5).

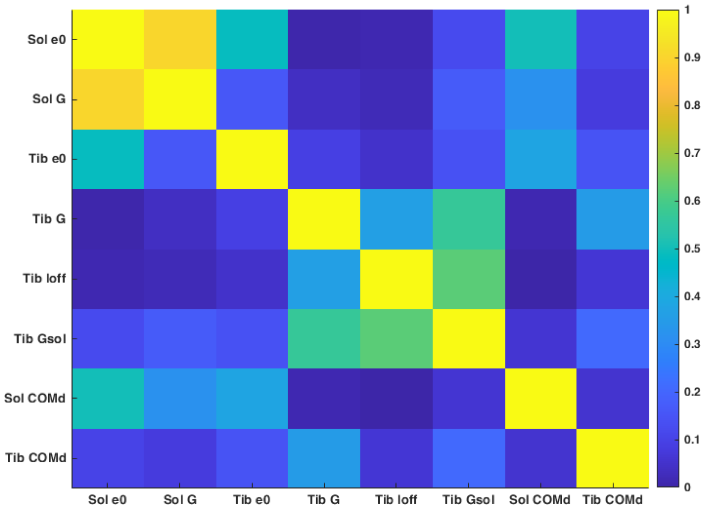

Figure S 5: **Covariance between control parameters**. We observed a strong (negative) correlation between the baseline soleus activity ($e_{sol,0}$) and the soleus force feedback gain ($G_{sol}$). Therefore, we decided to keep the baseline soleus activity constant ($e_{0,sol} = 0.027$) during the final parameter estimation process.

### 3 Supplementary information exoskelet experiment

#### 3.1 Adaptation to treadmill walking with exoskeleton

The experiment started with 20 minutes steady-state walking with the exoskeleton controlled with the default neuromuscular model to adapt to treadmill walking, with the heavy exoskeleton and the exoskeleton assistance. In addition to the decrease in muscle activity, we observed a gradual decrease in stride frequency and increase in mechanical work done by the exoskeleton during the adaptation (Fig. S7,6). It is however unclear if these changes during adaptation are mainly related to the adaptation to treadmill waking with the heavy exoskeleton or related to adaptation to the assistance.

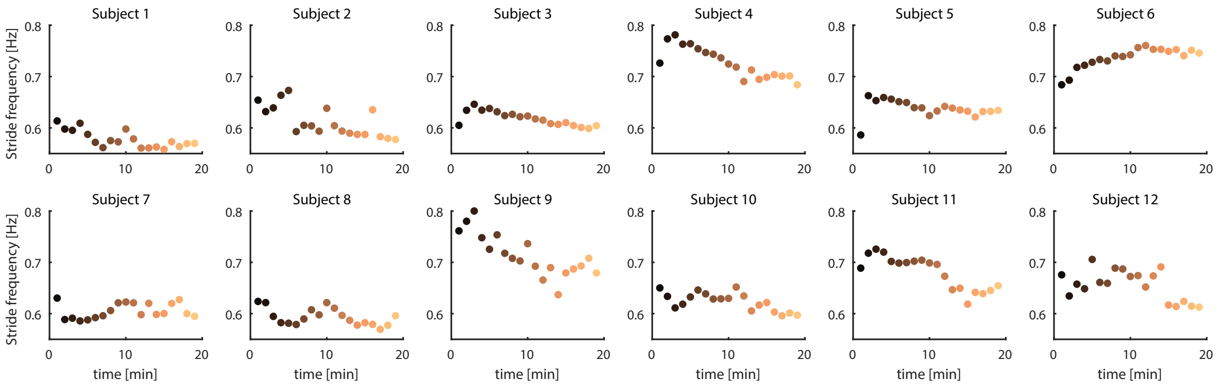

Figure S 6: 1 minute average stride frequency during the 20 minutes adaptation session.

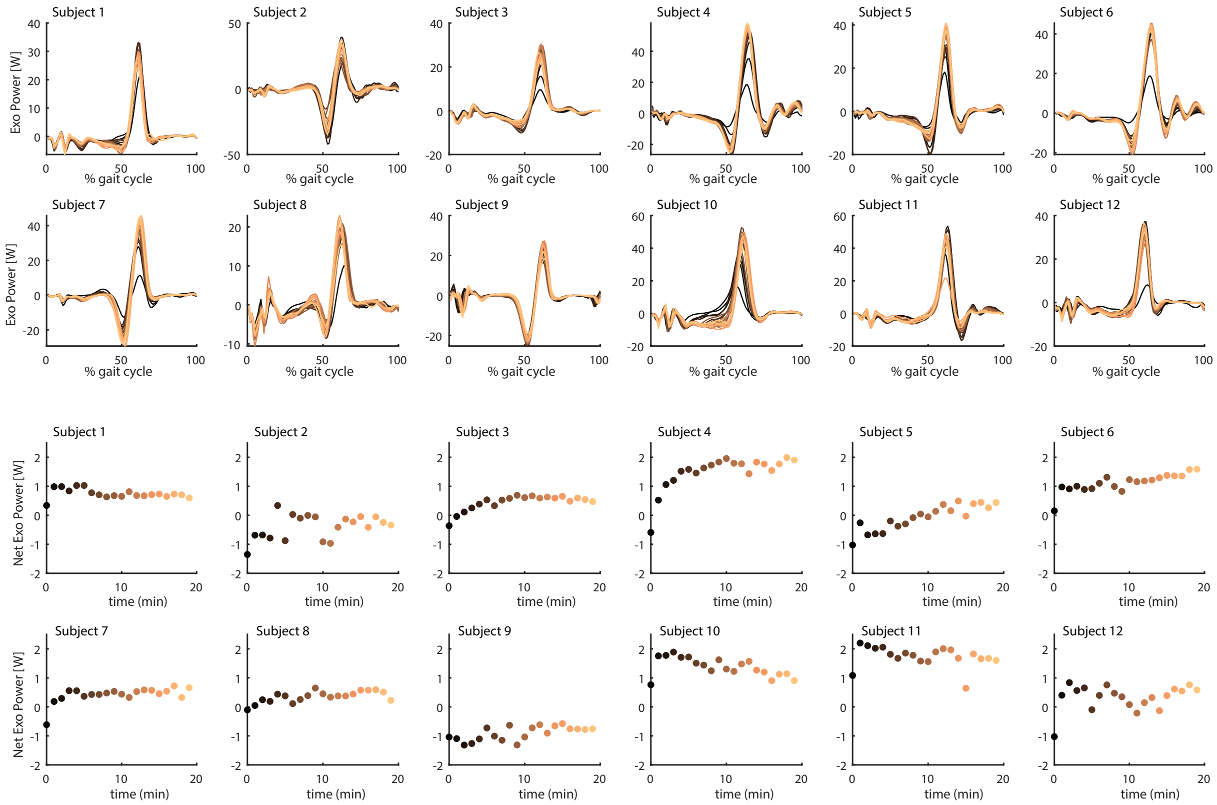

Figure S 7: Exoskeleton power as a function of the gait cycle and net exoskeleton power during the 20 minutes adaptation session.

#### 3.2 Muscle activity in steady-state walking

We used the unperturbed gait cycles during the time period in between perturbations to evaluate if both neuromuscular controllers reduce soleus activity during the perturbation session. Similar to the 19% reduction in at the end of the adaptation session, we observed a 20% reduction in soleus activity for both neuromuscular controllers during the unperturbed gait cycles between perturbations. This confirms that both neuromuscular controllers cause a similar reduction in soleus activity.

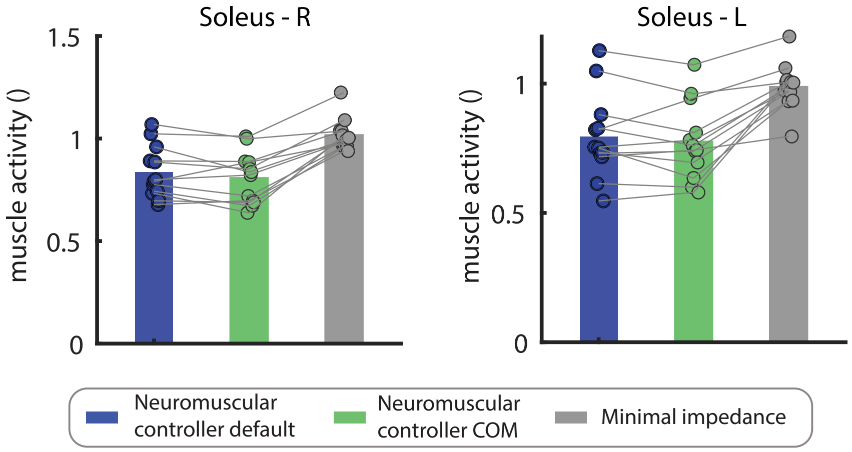

Figure S 8: **Average soleus activity during steady-state gait cycles in the last two minutes of the perturbation session.** Each dot represents the median soleus activity during the unperturbed gait cycles of an individual subject and the bars represent the average across subjects.

#### 3.3 Relation between COM kinematics and ankle moment in perturbed walking with exoskeleton

As an exploratory analysis we evaluated if the increase in muscle activity after perturbation is related the deviations in center of mass movement. This analysis is based on the observation that changes in muscle activity and the deviations in center of mass position and velocity are strongly correlated during perturbed walking without an exoskeleton (21).

We performed a very rudimentary analysis by correlating the COM displacement (displacement of the pelvis during the perturbed stance phase) and the average muscle activity during the first 500 ms after perturbations. Similar as in perturbed walking without an exoskeleton, we found that an increase in tibialis anterior activity is related to the backward movement of the center of mass and the increase in soleus activity is related to a forward movement of the center of mass (Fig S 9). This exploratory analysis provides a first indicition that subjects used a similar balance control strategy when walking with the exoskeleton compared to walking without an exoskeleton.

We believe that a new experiment with a range of perturbation magnitudes and longer adaptation times are needed to evaluate if human sensorimotor processing is altered by the exoskeleton support. Nevertheless, there is a reasonable variability in muscle activity and center of mass movement after the perturbation in our dataset, which enabled this exploratory analysis (Fig S 9. Note that this variability is similar as perturbed in walking without an exoskeleton. We believe that this variability is most likely related to variations state of the subjects when the perturbation was applied as we observed no clear adaptation effect during the perturbation session (Fig S 10).

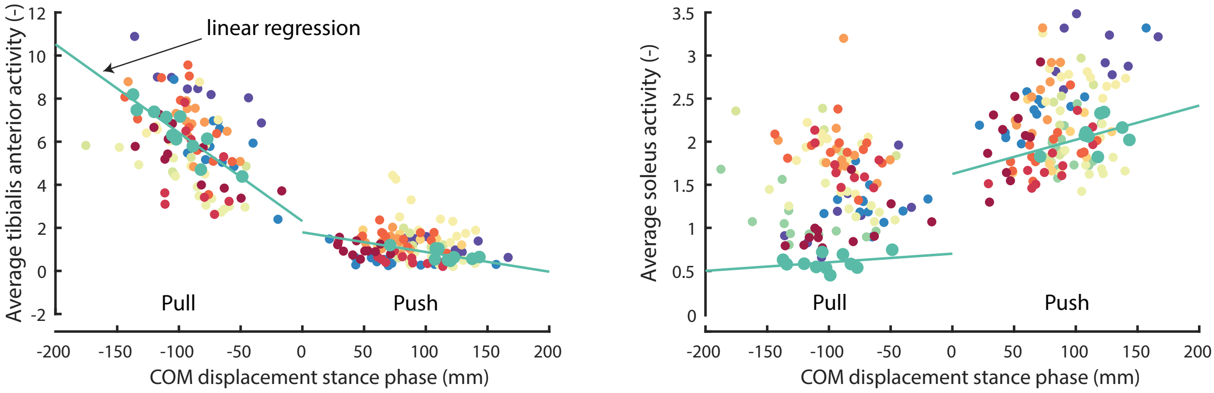

Figure S 9: **Average tibialis an soleus muscle activity and COM displacement in response to push and pull perturbation**. The color coding is used to identify the 12 subjects and each dot represents the response to single perturbation. This figure contains data of all the 12 subjects walking with the neuromuscular controller with COM velocity feedback. We highlighted one subjects with larger dots and a linear regression to highlight in a particular subject the relation between COM displacement and reactive muscle activity.

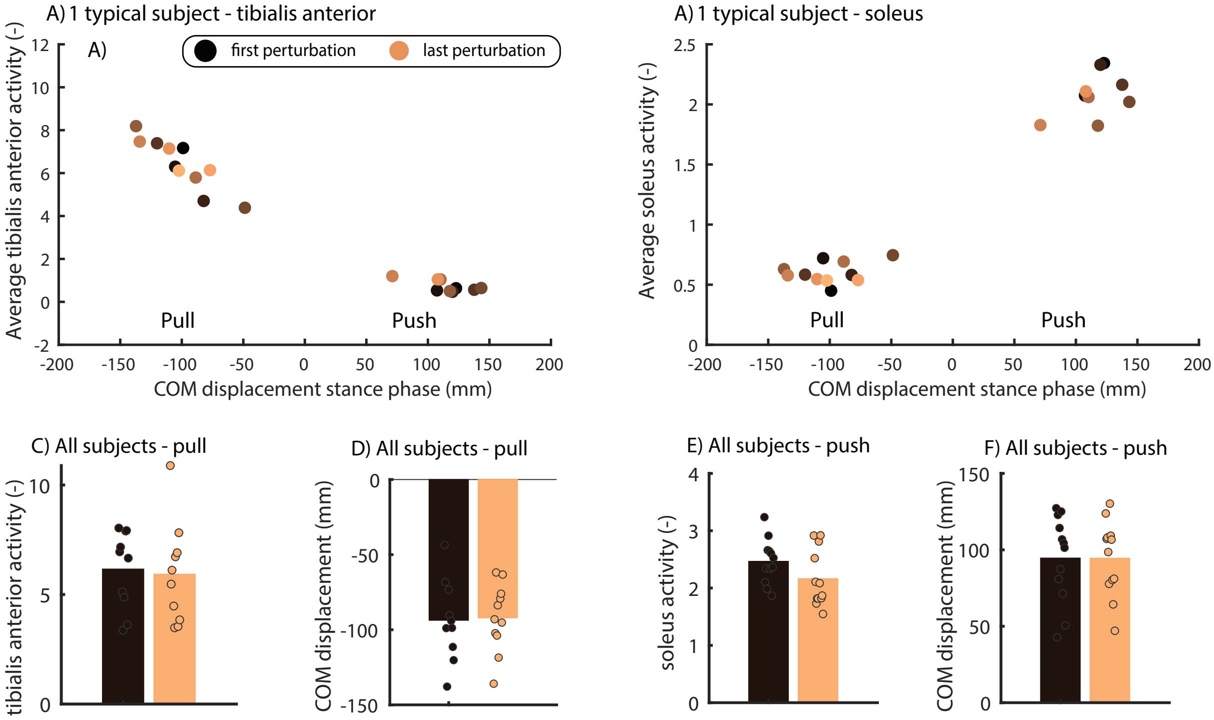

Figure S 10: **The response to the first and last perturbation in each session were similar**. The color coding shows changes in muscle activity and center of mass movement in response to the first (dark) and last (orange) perturbation in each session. A-B contains data of one typical subject. Bar plot C-F contain data of all the first (dark) and last (orange) perturbation in all subjects.
